# Transcutaneous cervical vagus nerve stimulation improves speech comprehension in noise: A crossover, placebo-controlled study

**DOI:** 10.1101/2024.12.10.627804

**Authors:** Michael Jigo, Jason B. Carmel, Qi Wang, Charles Rodenkirch

## Abstract

Speech comprehension in noisy environments remains a significant challenge, even among individuals with clinically normal hearing and users of hearing aids and cochlear implants. While conventional assistive hearing devices address limitations in the auditory periphery, they do not directly enhance the brain’s capacity to segregate speech from background noise. Because tonic vagus nerve stimulation (VNS) has demonstrated potential for rapidly improving central sensory processing, this study investigated whether tonic transcutaneous cervical VNS (tcVNS) can augment speech-in-noise intelligibility.

Two cohorts of older human adults (60-84 years) participated in a placebo-controlled, crossover study. Participants completed speech-in-noise assessments using either QuickSIN or AzBio sentences, while receiving tonic tcVNS to the neck, or placebo stimulation to the neck-shoulder junction. Speech-in-noise performance was assessed by measuring participants’ accuracy in repeating sentences presented at varying signal-to-noise ratios (SNR) within background babble.

Tonic tcVNS improved speech-in-noise intelligibility compared to placebo. At the group-level, the SNR threshold for 50% speech intelligibility (SNR-50) improved by 0.76 dB in QuickSIN (p=0.016) and by 0.38 dB in AzBio (p=0.045). For individual participants, 50% demonstrated improvements that met a minimum clinically important difference (MCID) of 1 dB. Tonic tcVNS evoked progressively greater improvements as SNR increased in QuickSIN (p=0.021) and AzBio (p=0.00023), with the largest gains above 0 dB SNR. In 55% of participants, tcVNS improved intelligibility beyond an MCID benchmark of 4.9% at 5 dB SNR. While the magnitude of tcVNS-evoked improvements was inversely related to baseline speech-in-noise impairment (p=0.028), with the most impaired individuals demonstrating the largest gains, it did not correlate with hearing loss severity (p=0.97) or age (p=0.88).

Our findings indicate that tonic tcVNS can evoke immediate and clinically meaningful enhancements in speech-in-noise comprehension. This suggests tcVNS may complement conventional assistive hearing technologies and inform novel therapies for sensory processing disorders.

## Introduction

Speech comprehension in noisy environments poses a significant challenge, especially for older adults and the 1.5 billion individuals worldwide with hearing loss^1,2^. Notably, 10-12% of people with clinically normal hearing still report difficulty hearing in noise^3,4^. Age-related declines in peripheral and central auditory systems exacerbate speech-in-noise (SIN) challenges in older adults^5,6^. While hearing aids, cochlear implants, and over-the-counter sound amplifiers can compensate for peripheral deficits, they do not resolve central limitations that continue to distort SIN perception^7–9^. Therefore, interventions targeting brain-based auditory processing could complement existing assistive hearing strategies to augment speech intelligibility in noise.

Vagus nerve stimulation (VNS) offers a promising method to address auditory processing limitations, with well-documented effects on the central nervous system. Invasive VNS, delivered via implanted electrode cuffs on the cervical branch in the neck, is an FDA-approved therapy for intractable epilepsy, pharmacological-resistant depression, and post-stroke motor rehabilitation^10,11^. More recently, transcutaneous VNS (tVNS), which stimulates the vagus nerve non-invasively via its cervical (tcVNS) or auricular (taVNS) branches, has emerged as a potential method for vagal activation^12–14^. The established safety and tolerance of tVNS^15^ in humans have spurred numerous studies in various applications including language learning^16–18^, cognitive performance^19,20^, and sensory processing^21–23^.

Recent preclinical^21^ and pilot clinical studies^23^ suggest that delivering VNS in a continuous, tonic fashion induces rapid and sustained central sensory improvements. Tonic VNS builds upon conventional paradigms, including duty-cycled protocols that deliver VNS for brief durations (e.g., 30 s ‘on’ followed by 60 s or more ‘off’^24–26^) and phasic protocols that pair VNS bursts with motor or sensory events to accelerate rehabilitation^11,27–29^. Though duty-cycled and phasic VNS effectively induce delayed, long-term neuroplasticity via short ‘on’ periods^21,22^, tonic VNS provides the distinct benefit of rapid, short-term sensory enhancements that are sustained through continuous vagal activation.

VNS-driven changes to sensory processing result from the activation of the locus coeruleus (LC)-norepinephrine (NE) system, which has long been hypothesized to contribute to the clinical benefits by VNS^30^. VNS engages the LC-NE system in both animals and humans^31–33^ through afferent vagal projections to the nucleus tractus solitarius^34^, which in turn sends excitatory signals to the LC via the nucleus paragigantocellularis^35^. In rodents, tonic electrical or optogenetic LC stimulation was shown to enhance sensory processing through its projections to the thalamus, a critical stage for sensory perception^36,37^. Tonic LC activation optimized thalamocortical neurons for sensory processing, increasing their efficiency and rate of sensory information transmission, and ultimately improving perceptual sensitivity during a go/no-go vibrotactile discrimination task^38^. Pharmacological manipulation revealed that improvements stemmed from a steady increase in NE concentration, which suppressed T-type calcium channel activity in the thalamus. This suppression reduced neural bursts that otherwise degrade sensory signal transmission^38^. Given the afferent vagal projection to the LC, tonic invasive VNS induced similar thalamocortical optimization that enhanced central sensory processing^21^.

Given the effects of tonic VNS on central sensory processing in rodents, a pilot translational study in humans explored the ability of tonic tcVNS or taVNS to improve sensory performance^23^. While tcVNS enhanced performance on visual and auditory psychophysics tasks, taVNS showed no such benefit. Crucially, tonic tcVNS improved performance in an auditory gap discrimination task^23^ that relied on the same precise temporal processing essential for speech perception^39,40^. Building on these findings, we hypothesized that tonic tcVNS could enhance speech-in-noise perception. To test this, we recruited older adults to receive tonic tcVNS or placebo stimulation while completing standard speech-in-noise assessments (QuickSIN^41^ and AzBio^42^) commonly used to guide hearing aid and cochlear implant recommendations^43,44^.

## Materials and Methods

### Ethics

Participants provided written informed consent and were compensated $30/hour. Protocols complied with the Declaration of Helsinki and were approved by Cornell University’s Institutional Review Board for Human Participant Research. Informed consent was provided for the images in **Figure 1** to be published.

**Figure 1.**
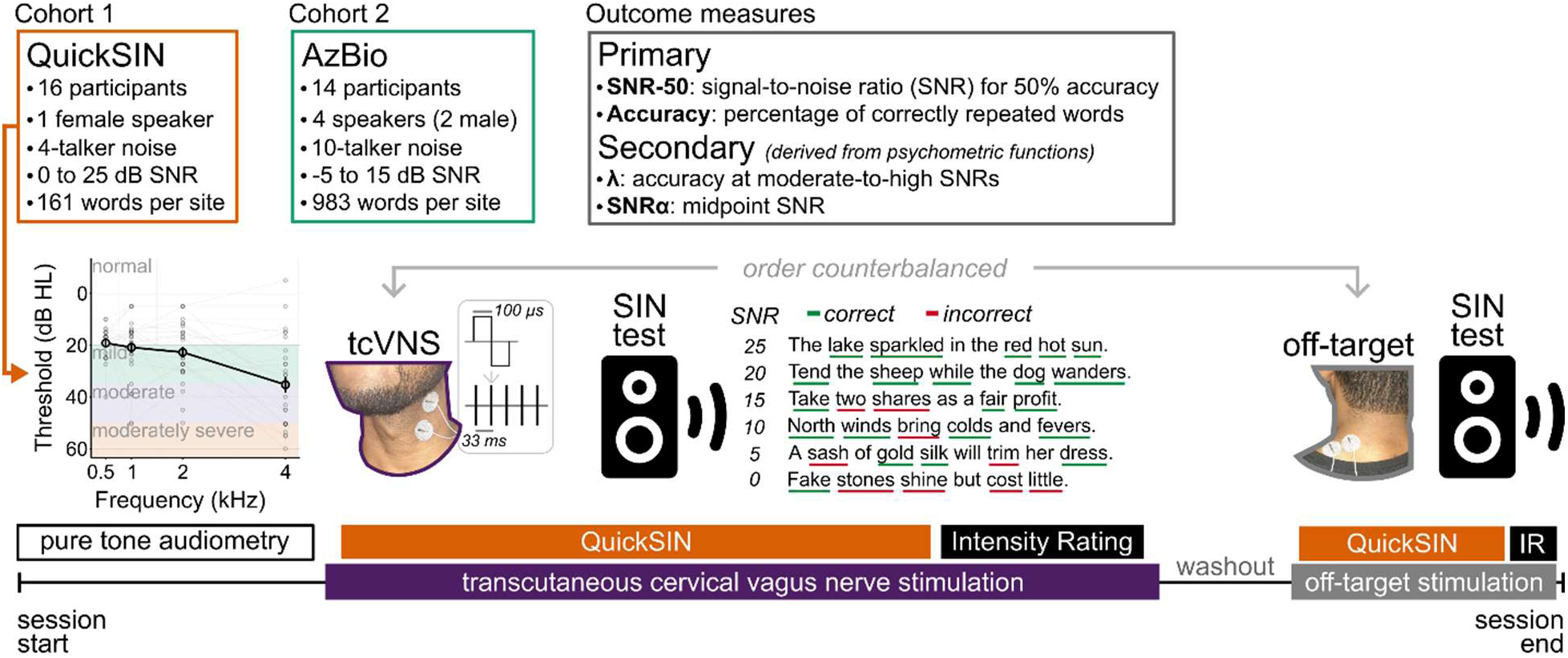
Speech-in-noise assessments and stimulation protocol. (***top***) Characteristics of each cohort, along with the study’s primary and secondary outcome measures. (***bottom***) Timeline showing events during a testing session for the QuickSIN cohort; an identical timeline was followed for the AzBio cohort. Pure tone audiometry was completed without electrical stimulation; data for all participants are shown, with large points and error bars showing the group-average and ±1 SEM connected by a dark black line. Hearing loss cutoffs are shown as different colors in the audiogram. Following pure tone audiometry, tonic tcVNS or off-target stimulation was delivered in a counterbalanced order; the stimulation waveform and frequency are shown beside ‘tcVNS’. After each speech-in-noise assessment, participants rated the perceived stimulation intensity.

### Protocol

We followed international standards for reporting the stimulation protocol and study design^45^, and summarize stimulation parameters in **Table 1**.

**Table 1.** Stimulation parameters and ratings of perceived intensity. Average values, 95% confidence intervals (within parentheses), and p-values for the difference between tcVNS and off-target parameters. Intensity ratings were measured with a 5-point Likert scale (1=not intense, 5=extremely intense).

|  |  | tcVNS | off-target | p-value |
| --- | --- | --- | --- | --- |
| QuickSIN | Current<br>mA | 7.71<br>(6.80, 8.62) | 8.15<br>(7.40, 8.90) | 0.13 |
|  | Intensity<br>1 = low, 5=high | 1.69<br>(1.44, 2.00) | 1.63<br>(1.19, 2.19) | 0.94 |
|  | Distance<br>cm | 3.84<br>(3.55, 4.12) | 3.04<br>(2.97, 3.11) | 0.00016 |
|  | Duration<br>min | 8.00<br>(7.29, 8.72) | 7.89<br>(7.23, 8.55) | 0.65 |
| AzBio | Current<br>mA | 8.59<br>(7.97, 9.21) | 8.86<br>(8.18, 9.55) | 0.25 |
|  | Intensity<br>1 = low, 5=high | 1.43<br>(1.07, 1.86) | 1.71<br>(1.29, 2.21) | 0.38 |
|  | Distance<br>cm | 4.00<br>(3.76, 4.24) | 3.04<br>(3.00, 3.09) | 0.0000024 |
|  | Duration<br>min | 20.95<br>(19.73, 22.17) | 20.93<br>(19.95, 21.91) | 0.95 |

### Stimulation sites and equipment

Hydrogel electrodes (diameter: 1”; PALS, Axelgaard Manufacturing, CA, USA) were placed on the skin, with center-to-center spacing adjusted for individual neck/shoulder anatomy. For tcVNS, electrodes were placed within the left carotid triangle—lateral to the larynx, medial to and oriented parallel to the sternocleidomastoid muscle. For off-target (placebo) stimulation, electrodes were placed over the trapezius muscle at the neck-shoulder junction where the vagus nerve is absent. Skin was prepped with hypoallergenic sanitary wipes.

### Study design

A placebo-controlled, single-blind, within-subject crossover design was used. Participants received tcVNS and off-target stimulation in a counterbalanced order, separated by a 29-minute washout period (95% CI: [25, 32]). Stimulation duration was determined by the duration of each test, which lasted approximately 8 minutes for QuickSIN and 21 minutes for AzBio cohorts, with sessions conducted in a single morning (9am-12pm) or afternoon (1pm-6pm).

### Stimulation parameters

An FDA-cleared neuromuscular stimulator operating in constant current mode (Mettler Sys*Stim 240, Mettler Electronics Corporation, CA, USA) delivered tonic, non-burst stimulation for both tcVNS and off-target conditions. Stimulation consisted of biphasic square pulses (100 μs/phase, 200 μs total) delivered at 30 Hz, in a continuous, tonic manner (100% duty cycle). Current amplitudes were individually calibrated using a standardized ramping procedure: current was initiated at 5 mA then increased gradually until muscle contractions or pain were reported, then decreased to the maximum level that was perceptible, tolerable, and did not induce muscle activation.

### Intensity rating

Immediately before stimulation ended, participants were prompted: “At this moment, how intense does the stimulation sensation feel?” and responded: 1=not intense, 2=slightly, 3=moderately, 4=very, 5=extremely intense.

### Blinding

The experimenter read a standardized cover story to participants: “The purpose of this study is to understand how electrical stimulation affects the clarity of your hearing. I will place a pair of electrodes at two different locations—on your neck or your upper back—that will activate the same nerve under your skin. While that’s happening, you will hear and repeat sentences played from a loudspeaker in the testing room.” Participants were not informed of the vagus nerve’s cervical branch location. Following this cover story, the experimenter described the QuickSIN or AzBio assessments. Electrical stimulation was delivered continuously during both ctVNS and placebo conditions to elicit similar physical sensations across conditions.

### Adverse Events

All adverse events were required to be reported to the IRB and to the medical oversight physician within 24 hours of occurrence. No adverse events occurred.

### Sample

#### Demographics

Participants were passively recruited from the Roosevelt Island community and Carter Burden Network. Inclusion criteria were: 60-85 years old, no history of neurological or psychiatric disorders, no history of substance abuse, not taking beta adrenergic blockers, no medical implants, no history of cardiac surgery, no severe cardiac disorders, no history of heart conduction abnormalities, not currently pregnant, no prior abnormalities or surgeries involving the neck or vagus nerve, and a passing score (≥12 of 15) on the 5-minute Montreal Cognitive Assessment (MoCA)^46^. Of the 43 individuals who inquired about the study, 19 declined to participate, and 2 were excluded due to a history of heart conduction abnormalities. The remaining 22 participants met eligibility criteria and provided informed consent. Testing occurred in two cohorts, 16 months apart, with eight individuals participating in both. The first cohort (n=16, 11 women; age=69.50 years; SD=7.87) was tested with QuickSIN^41^. Following advice from audiologists and an otolaryngologist, we conducted a conceptual replication of the QuickSIN findings in a second cohort with AzBio^42^ (n=14, 9 women; age=71.19 years; SD=6.70). Two QuickSIN participants wore hearing aids, and one AzBio participant was left-handed; all were included in the sample. Participants were asked to refrain from stimulants (e.g., caffeine, nicotine) for three hours prior.

#### Power analysis

While a sample size of 12 was sufficient to demonstrate statistically significant tcVNS-evoked sensory enhancements in younger adults^23^, we used a simulation-based approach^47^ to determine sample sizes in advance with ≥95% power for SIN perception. The smallest assumed change (SAC) we would observe was informed by studies deploying SIN assessments before and during sound amplification. For QuickSIN’s SNR-50, the SAC was 1.14 dB, the median improvement after two weeks of hearing aid use^48^. For AzBio, SAC for accuracy was 2.5%, based on improvements by a personal sound amplifier product^49^. To estimate the variance, six younger adults (5 women; age, 27.5 years; SD, 6.12) completed QuickSIN in a pilot that mimicked the present study in design and analyses. After 10,000 Monte Carlo simulations and assuming a 5% false positive rate, a sample of 16 had 95% power to detect the SAC for SNR-50 in QuickSIN, and 14 had 97% power to detect the SAC for SIN accuracy in AzBio.

### Speech-in-noise measures

QuickSIN and AzBio both involve repeating sentences read by a speaker, but AzBio is notably more complex, with varied voices, denser background babble, and more scored words (**Table 2**).

**Table 2.**
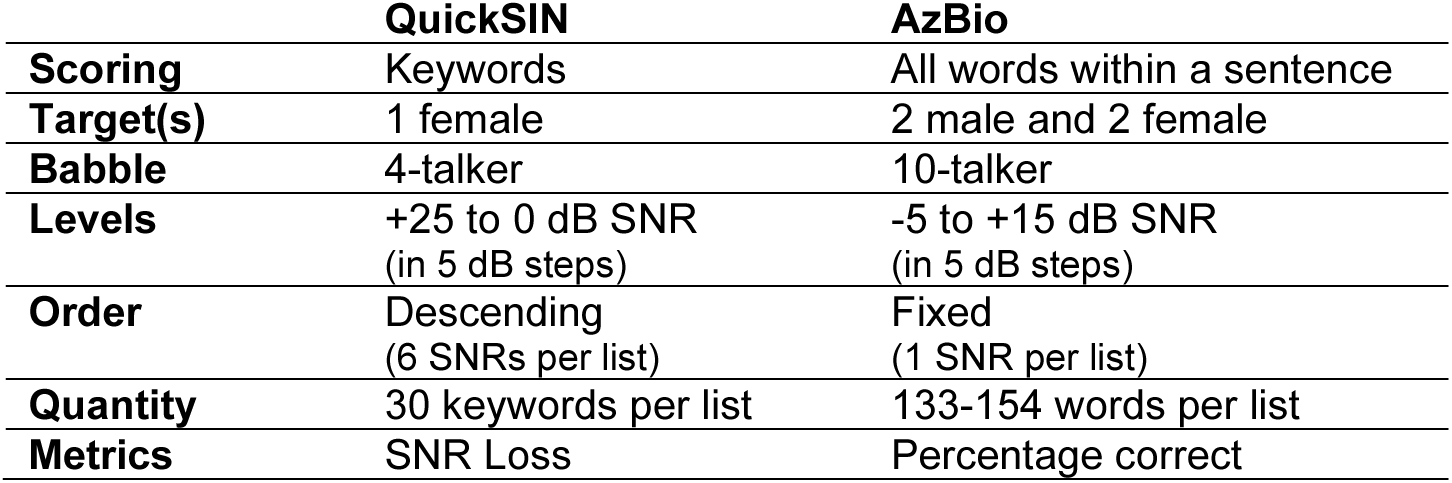
Speech-in-noise measures.

#### QuickSIN

Participants completed one practice list of sentences without electrical stimulation. Then, a median of 6 unique lists (30 keywords each) were played per stimulation site in randomized order. For each site, three participants completed 4 lists (8 total lists), 4 participants completed 5 lists (10 total), and 9 participants completed 6 lists (12 total). In each list, a female speaker read 6 sentences with increasing four-talker babble that decreased SNR from +25 to 0 dB in 5 dB decrements. Target speech was presented at 70 dBA SPL, adhering to guidelines^50^.

#### AzBio

We used AzBio Ver. 2.0^51^, featuring 15 equivalent lists (20 sentences each) spoken by two male and two female talkers against a 10-talker babble background (-5 to 15 dB SNR, 5 dB steps). Speech was presented at 60 dBA SPL, consistent with test guidelines^52^, except for one participant with 50 dB HL hearing sensitivity, who required 70 dBA SPL due to reported inaudibility. List 15 was used for practice and the remaining lists were ordered using a balanced Latin square design to control for first-order carryover effects. SNRs were fixed within and randomized between lists. Recommended SNRs (5, 10 dB)^44^ were tested twice, totaling seven lists per stimulation site.

#### Accuracy, SNR-50, and SNR Loss

SIN intelligibility was measured as the percentage of words accurately repeated. Following standard audiological testing procedures^53^, we estimated SNR-50 using Spearman-Kärber equations. QuickSIN is designed to use a simplified equation^50^ whereas for AzBio, we used a generalized form^54^. In QuickSIN, SNR-50 is typically converted to SNR Loss (SNR Loss = SNR-50 – 2); however, because the AzBio test does not have a similar conversion, we only report SNR-50.

#### Psychometric function

Three-parameter logistic functions were fit to accuracy as a function of SNR using quickpsy^55^: *p(SNR)=(1–λ) / (1+exp( -β[SNR–SNRα])*. SNRα defined its midpoint at low SNRs, λ defined its upper asymptote at high SNRs, and β defined its steepness. Pearson’s χ² determined goodness-of-fit.

### Apparatus

#### Sound field

Participants were tested in a sound-treated room. Stimuli were delivered via a single loudspeaker (JBL 308P MkII) using Psychtoolbox-3 in MATLAB on Windows 10, with audio processed through a digital-to-analogue converter and amplifier (Atom DAC+ and Amp+, JDS Labs). The system was calibrated for a flat frequency response (0.25-8 kHz) using a miniDSP UMIK-2 microphone and Room EQ Wizard (www.roomeqwizard.com). The loudspeaker was positioned 1 meter in front of participants (0° azimuth). Participant responses were streamed to an experimenter outside the room via a FIFINE K669B microphone and VB-Audio VoiceMeeter.

#### Audiometry

Monaural thresholds at 0.5, 1, 2, and 4 kHz were measured using a MAICO MA 40 audiometer and the modified Hughson-Westlake procedure, with the average in the better ear (PTA4) defining hearing loss. For QuickSIN participants, binaural thresholds (PTA-Binaural) were initially measured in the sound field using a single-interval, yes-no detection task and the QUEST^56^ method, converging on 50% hit rates. To estimate PTA4 from PTA-Binaural, seven QuickSIN participants completed a separate pure tone audiometry session with the audiometer. Their PTA4 and PTA-Binaural thresholds were highly correlated (r=0.91, p=0.0049), and we used linear regression (R^2^=0.82) to estimate PTA4 for all QuickSIN participants with the equation: *PTA4 = 13.67 + 1.05(PTA-Binaural)*.

### Statistics

All analyses were performed in R 4.4.0^57^.

#### Hierarchical modeling

Linear mixed-effects models (LME) were implemented in lmerTest^58^ and modeled differences between stimulation sites and signal-to-noise ratios as fixed effects. The variance among participants, lists, and/or SIN tests were random effects. Generalized LMEs with a binomial link function modeled changes in the percentage of correct responses. Bootstrapping estimated confidence intervals. P-values for fixed effects were determined with Satterthwaite approximations, following best-practice guidelines^59^, then adjusted for multiple comparisons to control the false discovery rate^60^. Post-hoc comparisons on marginal means were Bonferroni-corrected, as implemented in emmeans^61^.

#### Model Selection

Final LME models were selected from among candidates using the Bayesian Information Criterion (BIC) to balance model fit and complexity, favoring parsimonious models best supported by the data^62,63^. Models that produced a singular fit or could not converge were excluded. Details on model formulas are in **Supplementary Methods and Tables**.

#### Controlling for collinearity and regression to the mean

Multivariable LME models assessed the relationship between tcVNS effects and individual differences in baseline SIN performance (SNR-50 during placebo), hearing loss (PTA4), or age. All predictors were standardized to z-scores. tcVNS effects were defined as the difference in the upper asymptote of intelligibility between tcVNS and off-target conditions. Additional models included baseline SIN accuracy and MoCA scores as predictors. Variance Inflation Factors were used to assess multicollinearity. Permutation analyses, with maxT correction for multiple comparisons^64^, corrected for regression to the mean in the final model^65^.

#### Sex

Due to the small sample of each sex, and because sex-based differences were not the primary focus of this study, sex-related effects were assessed in exploratory and post-hoc analyses. SNR-50 was pooled across cohorts to create a larger sample of males, then LMEs evaluated whether tcVNS efficacy interacted with sex.

#### MoCA, PTA, and Stimulation Parameters

Exploratory, post-hoc analyses tested differences between cohorts’ MoCA scores and PTA thresholds (Kruskal-Wallis test), differences in stimulation parameters (paired t-test), and differences in perceived stimulation intensity (paired Wilcoxon signed rank test).

## Results

### Tonic tcVNS was well-tolerated

Before speech-in-noise (SIN) testing, participants in two cohorts underwent cognitive screening (MoCA) and pure tone audiometry (PTA) without electrical stimulation (**Figure 1**). All participants passed the cognitive screen (≥12/15 points), but MoCA scores differed between cohorts (H=4.38, p=0.030), with the QuickSIN cohort showing higher scores (mean=13.5, SD=1.15) than the AzBio cohort (mean=12.6, SD=0.84). PTA thresholds (PTA4) did not differ significantly between cohorts (H=1.17, p=0.28), with both exhibiting mild hearing loss on average (QuickSIN: mean=26 dB HL, SD=8.22, range=[15, 40]; AzBio: mean=23 dB HL, SD=11.69, range=[5, 50]).

During SIN testing, each participant received both tonic tcVNS and off-target (placebo) stimulation in a counterbalanced order, which minimized order effects (see **Supplementary Analyses**). Continuous tonic stimulation (biphasic square pulses delivered at 30 Hz) was well-tolerated, with its perceived intensity rated between “not intense” and “slightly intense” on average (**Table 1**).

### Speech intelligibility thresholds improved during tcVNS

SNR-50, the signal-to-noise ratio required for 50% speech intelligibility, was estimated using the Spearman-Kärber method^54^, with lower values indicating better SIN performance. Linear mixed-effects models (; **Table S1**) found a significant reduction in SNR-50 during tcVNS compared to off-target stimulation: 0.76 dB in QuickSIN (**Figure 2A**; SE=0.30, t(155.03)=-2.53, p=0.016) and by 0.38 dB in AzBio (**Figure 2B**; SE=0.17, t(13)=-2.22, p=0.045**; Table S2**), corresponding to improvements of 12% and 13%, respectively. Exploratory analyses found no evidence of sex-related differences in both cohorts (**Table S3-S4**; interaction: t(174.98)=0.094, p=0.93).

**Figure 2.**
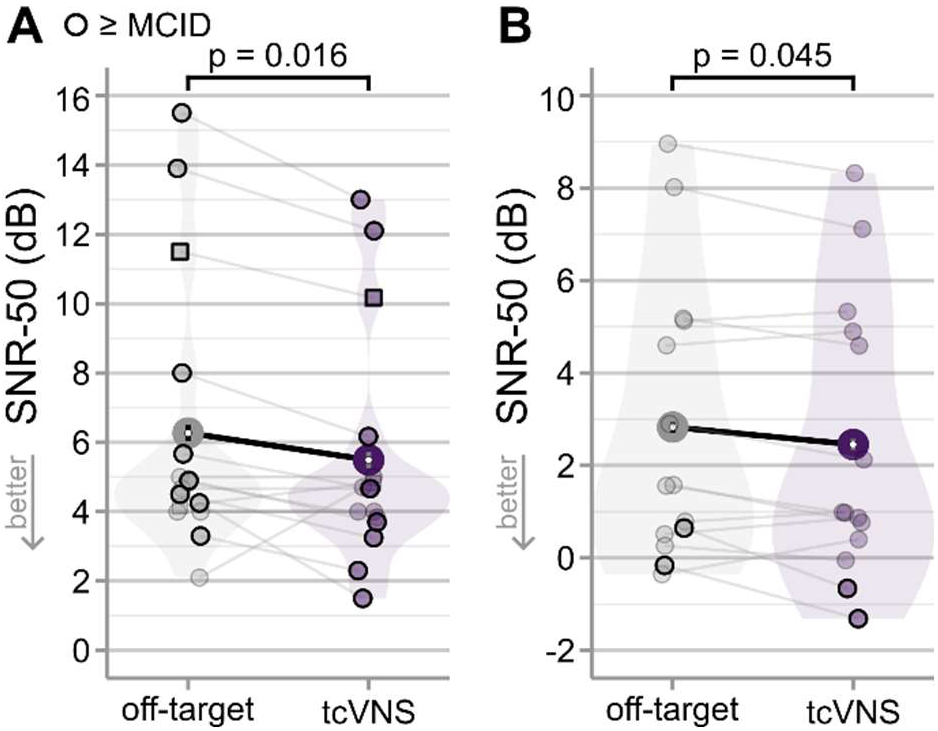
Speech intelligibility thresholds improved during tcVNS. SNR-50 estimated with (**A**) QuickSIN and (**B**) AzBio. Lower values indicate better speech-in-noise performance. Two individuals in the QuickSIN cohort wore hearing aids during the test and are depicted as square markers. Participants in which tcVNS improvements met or exceeded the minimum clinically important difference (MCID) are highlighted with darkened outlines. Large data points connected by black lines show group means, with errors ±1 within-subject SEM^67^. P-values are corrected for multiple comparisons using the Benjamini-Hochberg procedure.

Improvements by tcVNS met the 1 dB minimum clinically important difference (MCID) established for an FDA-approved cochlear implant (Nucleus 24^66^) in 9 of 16 QuickSIN participants (56%; **Table 3**) and in 2 of 14 AzBio participants (14%). Overall, 11 of the 22 individuals in the study (50%) demonstrated MCID-level improvements, suggesting a clinically meaningful benefit. This overall count reflects all individuals in the study, with those participating in both cohorts counted only once if they met the MCID in either test. Therefore, this total is not the sum of the two cohort results.

**Table 3.**
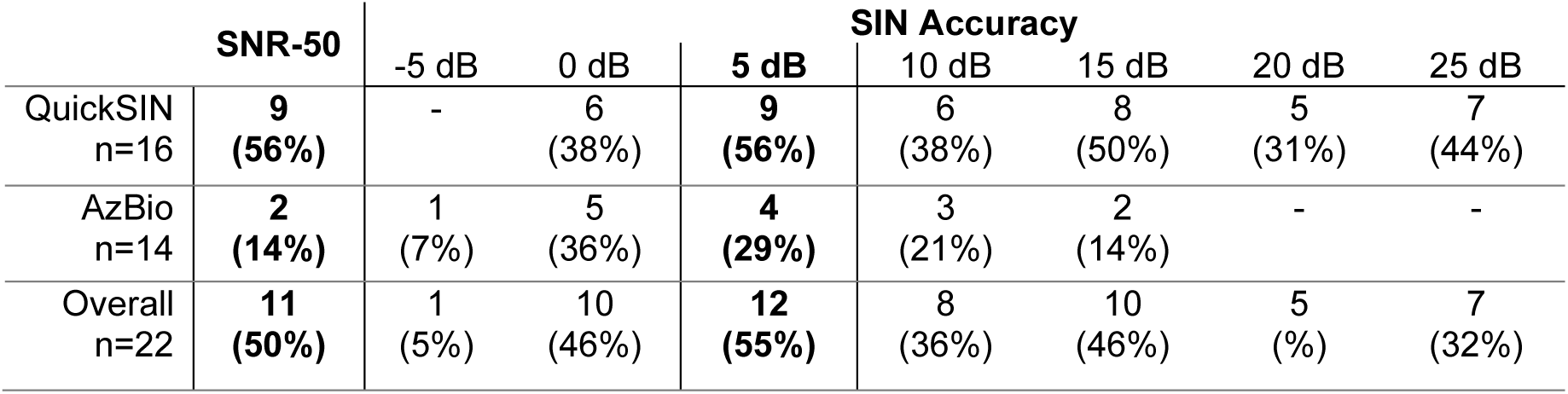
Improvements to speech-in-noise performance met established minimum clinically important difference (MCID) benchmarks. Displayed are the number and percentage of participants in which improvements by tcVNS met or exceeded the MCID for SNR-50 (≥1 dB) or word recognition accuracy (≥4.9%). The “Overall” row represents the total number of individuals who demonstrated MCID-level improvements across both cohorts. Individuals that participated in both cohorts are counted once if their performance met the MCID in either test. As a result, the “Overall” row may not equal the sum of each individual cohort.

### tcVNS improved speech intelligibility in low-to-moderate noise

The relationship between SIN intelligibility and SNR was well-fit by logistic functions (**Figure 3**; χ²>12, p>0.067). This function is defined by two key parameters^68^: SNRα, the signal-to-noise ratio at its midpoint, and λ, the upper asymptote of accuracy at high SNRs. LME models (**Table S5-S6**) indicated that tcVNS increased the upper asymptote by 3.74% in QuickSIN (**Figure 3A**; SE=1.43, t(15)=2.62, p=0.031) and by 2.43% in AzBio (**Figure 3B**; SE=0.99, t(13)=2.45, p=0.039). However, tcVNS did not significantly change the function’s midpoint (QuickSIN: t(15)=-0.066, p=0.95; AzBio: t(13)=0.15, p=0.95), suggesting its benefits are primarily on SIN intelligibility in low-to-moderate noise.

**Figure 3.**
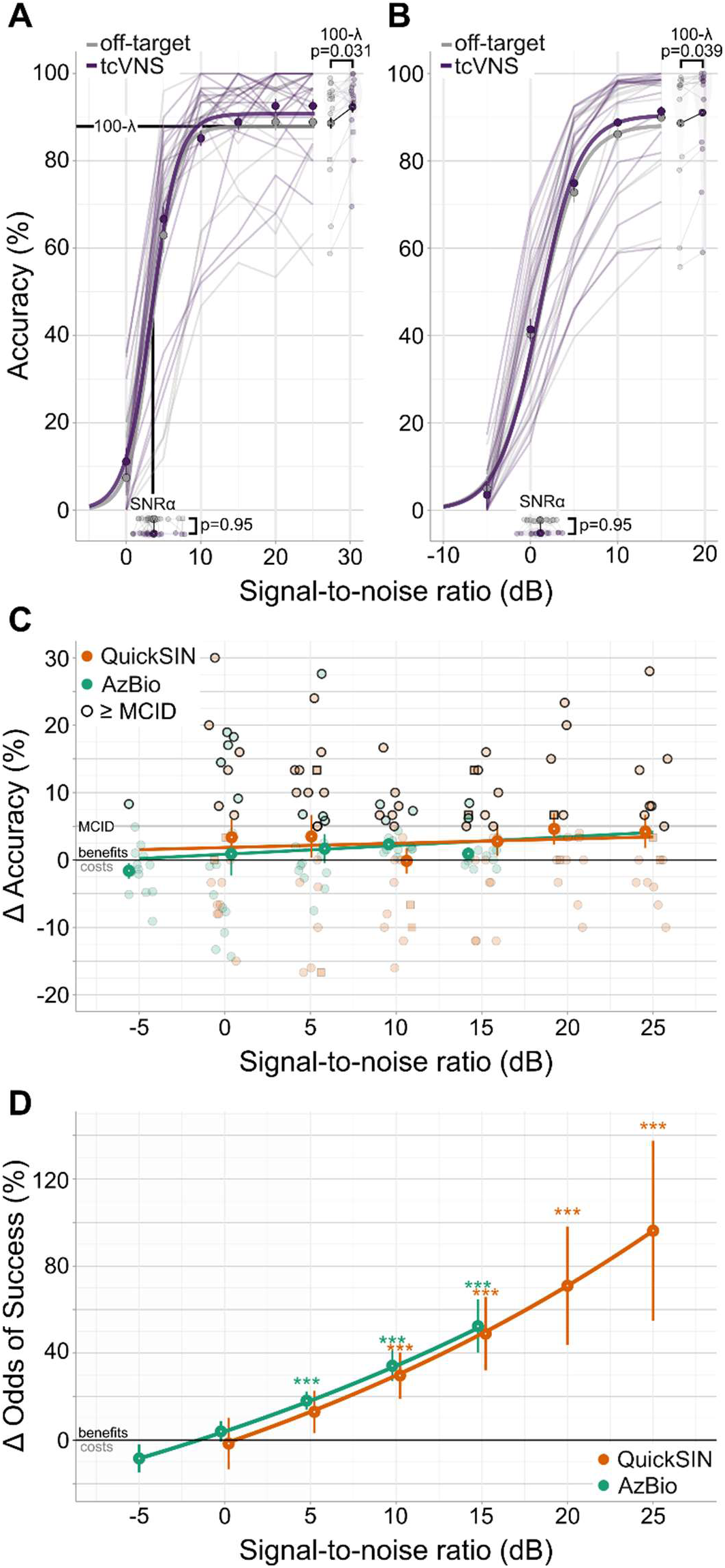
tcVNS improved speech intelligibility in low-to-moderate noise. Psychometric functions for (**A**) QuickSIN and (**B**) AzBio. Smooth lines depict the fitted logistic functions while the transparent, jagged lines show the raw accuracy for individual participants. Parameter estimates for SNRα and λ are displayed beside the x- and y-axes, respectively, with large points connected by black lines indicating the group average and error bars of ±1 within-subject SEM^67^. P-values correspond to the result of linear mixed-effects modeling and are corrected for multiple comparisons using the Benjamini-Hochberg procedure. (**C**) Change in SIN intelligibility as a function of SNR. Positive values indicate better accuracy during tcVNS. Large points and errors bars show the group-average difference and ±1 within-subject SEM^67^; smaller points show individual participants, with those meeting the minimum clinically important difference (MCID) highlighted with darkened outlines. The best fitting regression lines for each cohort are displayed. (**D**) Percent change in the odds of accurate SIN comprehension. Positive values indicate higher odds during tcVNS. Points and vertical lines depict the marginal means and their standard error. Star signifiers denote Bonferroni-corrected p-values for post-hoc comparisons: ***: p<0.001.

Generalized LME models (**Table S7-S8**) demonstrated that tcVNS-evoked improvements increased with SNR (**Figure 4C**), with a 0.028 log odds increase per SNR in QuickSIN (SE=0.011, z=2.42, p=0.021) and by 0.025 log odds in AzBio (SE=0.0067, z=3.81, p=0.00023). Post-hoc comparisons of marginal means found significant performance gains at SNRs above 0 dB. These comparisons were conducted on the odds ratio scale, which is well-suited for evaluating the binary outcome of correct versus incorrect SIN perception. In QuickSIN, tcVNS improved the odds of accurate SIN comprehension by 43% on average (**Figure 4D**), with significant effects at 10 (z=3.15, p=0.0098), 15 (z=3.52, p=0.0026), 20 (z=3.37, p=0.0046), and 25-dB SNR (z=3.20, p=0.0083). In AzBio, the odds of accurate performance increased by 20%, on average, with significant improvements at 5 (z=4.79, p=8.49×10^-6^), 10 (z=5.67, p=7.10×10^-8^), and 15-dB SNR (z=5.27, p=7.00×10^-7^).

**Figure 4.**
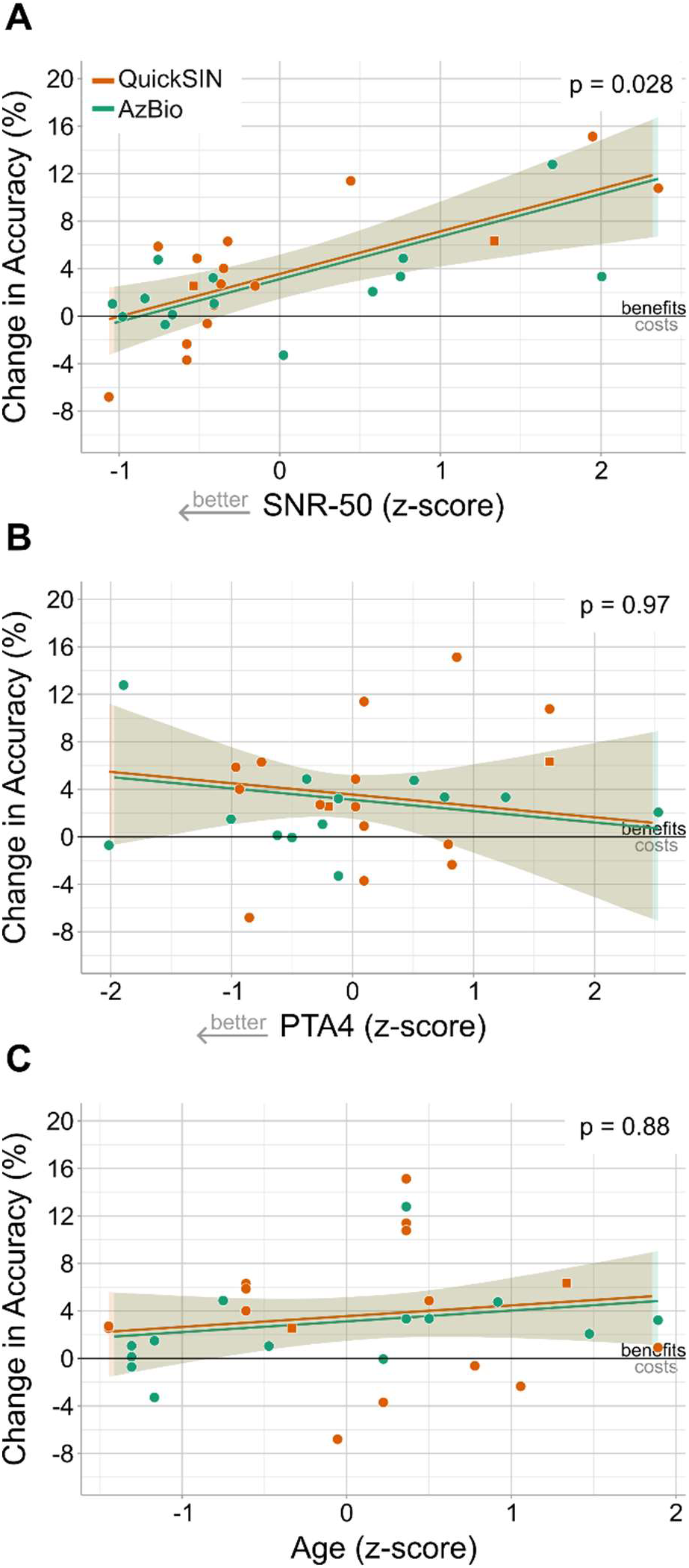
Improvements grew with the severity of SIN deficits, but neither hearing loss nor age. tcVNS-evoked changes in the upper asymptote of SIN accuracy (λ), are displayed as function of (**A**) baseline SNR-50 measured during off-target stimulation (**B**) hearing loss (PTA4), and (**D**) age. In all panels, each dot depicts individual participants in a cohort, lines depict the marginal regression for each independent variable determined using LME modeling, with shaded areas indicating 95% confidence intervals. All p-values were corrected for regression to the mean and for multiple comparisons using the maxT method^64^.

Individual participants showed clinically significant gains (**Table 3**). A 4.9% improvement to SIN intelligibility at 5 dB SNR defined the MCID^69^. While average tcVNS-evoked improvements at 5 dB were 3.02% in QuickSIN (SE=3.22) and 2.33% in AzBio (SE=2.23), 56% (9/16) of QuickSIN and 29% (4/14) of AzBio participants met the MCID. Overall, 55% (12/22) demonstrated MCID-level improvements, supporting the potential for meaningful clinical benefits with tcVNS.

### Improvements grew with the severity of SIN deficits, but neither hearing loss nor age

To explore the variability in individual responses to tcVNS, we analyzed whether differences in age (mean=70.4 years, SD=7.19), hearing loss, or baseline SIN performance influenced efficacy. SNR-50 scores during off-target stimulation indexed baseline SIN ability (QuickSIN: mean=6.27 dB, SD=3.92; AzBio: mean=2.83 dB, SD=3.06), and pure tone thresholds (PTA4) indexed hearing loss.

Participants exhibited varying degrees of SIN deficits and hearing loss. Using normative SNR-50 data for QuickSIN^50^, 69% of participants had normal/near-normal SIN performance (<5 dB SNR-50), 12% were mildly impaired (5 to 9 dB), and 19% moderately impaired (9-17 dB). Although equivalent categories are less established for AzBio, a higher SNR-50 indicated greater SIN deficits. Hearing loss also varied: 27% had normal hearing (<20 dB HL), 60% had mild (20 to <35 dB HL), 10% had moderate (35 to <50 dB HL), and 3% had moderately severe hearing loss (50 to <65 dB HL). As expected, SIN deficits correlated with hearing loss and age (**Figure S1** and **Tables S9-S10**).

Exploratory LME modeling evaluated whether these individual differences influenced tcVNS efficacy. Baseline measures of SNR-50, PTA4, and Age were included as predictors, while other baseline measures (MoCA scores and SIN accuracy), were excluded due to high collinearity (Variance Inflation Factors >6.7; **Table S11-S12**). Given that tcVNS had the largest effects at moderate-to-high SNRs, changes in the upper asymptote (λ) quantified its efficacy. Results showed that tcVNS-evoked improvements grew by 3.59% (SE=1.01, t(21.66)=3.55, p=0.028) for each standardized unit of change in baseline SNR-50 (**Figure 4A**). However, neither hearing loss (t(21.03)=-0.63, p=0.97; **Figure 4B**), age (t(21.16)=0.89, p=0.88; **Figure 4C**) nor any two-way or three-way interactions significantly predicted tcVNS-evoked improvements (p>0.93; **Table S13**). These findings suggest that tcVNS efficacy was primarily determined by the severity of pre-existing SIN deficits rather than hearing loss or age.

## Discussion

Age-related declines in auditory processing impair speech comprehension in noisy environments. Across two cohorts of older adults performing standard SIN assessments, tonic tcVNS improved clinically relevant metrics used to guide hearing aid and cochlear implant recommendations^43,44^. SNR-50 thresholds improved by an average of 0.76 dB in QuickSIN and 0.38 dB in AzBio. Speech-in-noise intelligibility increased by 3% when speech was louder than noise (SNRs >0 dB), with larger gains for individuals with pre-existing SIN deficits. Overall, tcVNS met the MCID for SNR-50 in 50% of participants and for SIN intelligibility in 55%, suggesting that tcVNS may meaningfully alleviate difficulties understanding speech in noise.

Given these significant improvements in SIN performance, we explored how tonic tcVNS measures against established solutions: hearing aids, personal sound amplification products (PSAP), and cochlear implants. For instance, two weeks of use with self-fitted or audiologist-fitted hearing aids (Lexie Lumen) yielded SNR-50 improvements in QuickSIN of 0.47 dB and 1.14 dB, respectively^48^. Here, eight minutes of tonic tcVNS produced a comparable 0.76 dB improvement, placing its gains in-between these hearing aid results. In an assessment of five PSAPs, intelligibility with AzBio sentences improved by an average of 5.8%, though with considerable variation ranging from -11% to +11%. The 3% improvement observed during tonic tcVNS fell well within this range, near the average PSAP benefit. However, the 12% improvement reported for a hearing aid (Oticon Nera) exceeded tcVNS^49^. It is important to note that the PSAPs and hearing aid were tested with a 180° spatial separation between speech and noise, providing an additional cue to aid in speech perception^70,71^. Our study had speech and noise co-located at 0°, eliminating spatial cues. Moreover, while our cohorts included individuals with clinically normal hearing, they are typically not the target population for studies on assistive hearing devices^44,48,49^. Therefore, even when tested under more challenging conditions and with a wider range of hearing sensitivities, tonic tcVNS paralleled conventional assistive hearing solutions, with MCID-level gains in QuickSIN^66^ and AzBio^69^.

Our findings also suggest that tonic tcVNS may improve SIN intelligibility in realistic communication scenarios. During normal conversation, speakers adjust their speech levels to overcome background noise^72–75^. As a result, speech is usually louder than background noise by 5-10 dB in everyday settings including stores, restaurants, and in moving traffic^73–75^. Notably, improvements by tonic tcVNS grew with SNR, delivering significant gains at SNRs ≥5 dB, which implies its effects are well-adapted to the demands of real-world listening situations.

Growing evidence support the effectiveness of tonic tcVNS for enhancing central auditory processing^21–23,38^. Sensory processing depends on neuromodulatory systems that regulate attention and arousal, such as the LC-NE^22,76^. In line with this, tonic LC-NE activation, either directly^38^ or indirectly via VNS^21^, has been shown to rapidly improve and sustain accurate central sensory processing. In rodents, tonic LC stimulation resulted in a steady elevation of NE concentration, which suppressed burst spiking responses that would otherwise reduce the rate and efficiency of sensory transmission^21,38^. While direct evidence of this suppressive mechanism in humans is lacking, our prior work in younger adults suggests that tonic tcVNS improves sensory performance in a manner consistent with this mechanism of action^23^. Specifically, tonic tcVNS improved the detection of brief gaps in sound, a task that depends on precise temporal processing by the central auditory system^39,40,77^ that declines with age and limits speech intelligibility^40,77^. Therefore, our present results provide support for tcVNS-induced gains to central auditory processing that facilitate accurate speech perception in young and old adults.

The improvements observed during tonic tcVNS may also derive from gains in selective attention and working memory—key cognitive functions for successful SIN perception^78–81^. The LC-NE system has been shown to modulate goal-directed attentional selection^76,82^ and maintain stable working memory capacity^83,84^, shaping cognitive function in aging^85^. In older adults, deficits in these functions are exacerbated by age-related hearing loss, which accelerates cognitive decline^86,87^. Impaired attention limits the ability to focus on relevant speech while suppressing background noise, while working memory deficits reduce the retention of recently heard words, increasing susceptibility to distraction in multi-speaker environments^88,89^ and weakening the utility of contextual cues^6,90^. Deficits in these functions have been linked to a disrupted excitation-inhibition balance within thalamocortical circuits that support rapid temporal processing and auditory perceptual organization^91^. Increased LC activation, and a concomitant elevation in NE concentration, has been shown to increase the efficacy of excitatory (glutamic) and inhibitory (GABAergic) responses within the thalamus^38,92^, providing a potential mechanism by which tcVNS-mediated activation of the LC-NE could alleviate excitation-inhibition imbalances to improve cognitive function for SIN perception.

Other neuromodulation approaches have shown limited success in improving SIN intelligibility. Transcranial direct current stimulation (tDCS) over the left superior temporal gyrus improved SIN performance in the CUNY Sentences Test^93^, but used a sham condition without adequate control for the tDCS sensation^94^. In contrast, tonic tcVNS and placebo stimulation were perceived as equivalently intense in our study. Intermittent theta-burst transcranial magnetic stimulation was initially found to improve SIN intelligibility when delivered to the left ventral premotor cortex^95^. However, a follow-up study did not replicate these effects^96^. Instead, benefits occurred only in individuals with poor baseline performance, like the compensatory benefits of tonic tcVNS we found in younger adults^23^ and, now, in older adults. Lastly, pairing brief VNS bursts with auditory tones or sensory training can accelerate and strengthen recovery from sensory disruption^11,27–29^. However, in a rat model of hearing loss, pairing VNS with training did not improve speech perception beyond training alone^97^. This contrast between the SIN enhancements we observed in humans with and without hearing loss, and the lack thereof after phasic VNS in an animal model of hearing loss suggests that stimulation parameters play a critical role for effective neuromodulation.

Our stimulation protocol was designed using prior research. First, the off-target stimulation site has been shown to have no effect on sensory performance, matching the impact of forearm stimulation or a no-stimulation baseline in previous work^23^. Second, the electrode montage’s spatial configuration has demonstrated auditory enhancements in humans^23^ and is supported by computational modeling indicating that our parameters can drive action potentials in afferent and efferent fibers of the vagus nerve’s cervical branch^12^. Moreover, based on prior finite-element simulations^98,99^, we used a farther inter-electrode spacing during tcVNS to enhance electrical field penetration and improve the likelihood of vagal activation. Third, our tcVNS protocol has been shown to modulate heart rate variability^23^, a marker of efferent vagal activation^100^. Lastly, given that VNS-evoked thalamocortical optimization dissipates within seconds after stimulation ends^21^, our approximately 30-minute washout period ensured a return to baseline between tcVNS and placebo conditions—a shorter duration (20 minutes) yielded robust tcVNS effects in prior work^23^.

Our findings corroborate evidence that tcVNS can improve sensory and cognitive performance in humans^18–20,23,101–103^, with its effects exceeding those of taVNS when compared head-to-head^18,23^. Although we ascribe our findings to a tcVNS-mediated activation of the LC-NE system, evidence supporting that link is mixed. While functional magnetic resonance imaging studies have demonstrated evidence of tcVNS-evoked activity in afferent vagal projections to the LC^32,33^, data from non-invasive NE biomarkers, including pupil diameter, P300, and salivary alpha amylase are inconclusive^104^. This variability has been attributed to studies being statistically underpowered, a lack of an appropriate active placebo control, and differences in stimulation parameters^104^. Although our study had design elements that addressed some limitations—power analyses, conceptual replication in two SIN tests, an active placebo control, and stimulation parameters informed by prior studies—future work is required to replicate our findings and confirm the LC-NE system’s role.

Lastly, our study had several limitations. First, two participants in the QuickSIN cohort wore hearing aids during SIN testing, introducing a potential confound. However, repeating all statistical analyses without these participants did not alter the pattern of results, so both were included in the final dataset (marked with square symbols in all plots). For one hearing aid user, tcVNS evoked MCID-level gains in SNR-50 (**Figure 2A**) and recognition accuracy (**Figure 4C**), suggesting that tcVNS complemented their hearing aids to enhance SIN perception. Future studies are needed to explore this interaction. Second, we did not formally evaluate blinding integrity; therefore, it remains uncertain whether participants could guess the active condition. However, several factors likely minimized this possibility: the perceived intensity between tcVNS and placebo was matched; participants were read a standardized cover story with no mention of vagal anatomy; and participants were laypersons without expertise regarding the vagus nerve’s cervical branch. Third, because VNS-mediated LC-NE activation can increase arousal to regulate performance^19,76^, it is plausible that group-level improvements may have been driven primarily by individuals participating during their off-peak hours when arousal is low, typically in the afternoon/evening for older adults^105^. While our sample size was not powered to assess this, future work is needed to distinguish general arousal effects from sensory enhancements by tcVNS.

In conclusion, tonic tcVNS evoked immediate and clinically meaningful improvements to speech perception in noise, particularly for individuals with pre-existing speech-in-noise deficits. This study adds converging evidence for an LC-NE-mediated enhancement to auditory processing in the brain, triggered on demand by tonic tcVNS. Our findings highlight the potential for tcVNS to complement conventional assistive hearing technologies and offer promising implications for novel treatments of sensory disorders.

## Acknowledgements

We thank Dr. Frank R. Lin, MD, PhD, Dr. Nicholas S. Reed, Ph.D, and Dr. Joseph J. Montano, Ed.D for useful discussions and comments, and the Weill Cornell Medicine-Hearing and Speech Center for providing us with an audiometer. This work was supported by Sharper Sense, Inc. and The Jacobs Technion-Cornell Institute at Cornell Tech. Qi Wang was supported by the National Institutes of Health [R01NS119813, R01AG075114, R21MH125107], National Science Foundation [CBET 1847315, TI 2232149] and the Air Force Office of Scientific Research [FA9550-22-1-0337]. Any opinions, findings, and conclusions or recommendations expressed in this material are those of the authors and do not necessarily reflect the views of the United States Air Force.

## Author Contributions

M.J., J.B.C, Q.W., and C.R. conceived the study. M.J. led the design of the study, with input from J.B.C., Q.W., and C.R. M.J. led participant recruitment, data collection, and data analysis. M.J. drafted the article and figures, with input from J.B.C, Q.W., and C.R. All authors approved the final manuscript.

