## Supplementary Materials for "Transcutaneous cervical vagus nerve stimulation improves speech comprehension in noise: A crossover, placebo-controlled study"

### Supplementary Methods

**Model formulation.** Initial model formulas were based on the research hypothesis, the structure of each speech-in-noise test, and the repeated-measures experimental design. Specifically, the fixed effects of Site (2 levels: tcVNS and off-target), SNR (QuickSIN: 6 levels; AzBio: 5 levels), and their interaction were central to our research hypothesis and experimentally manipulated in the study. Other exploratory analyses assessing sex- and order-related interactions included Sex (2 levels) and Order (2 levels: placebo-first, tcVNS-first) as fixed effects. For the random effects, Subj accounted for the repeated-measures design, Test accounted for variability due to differences between speech-in-noise tests (see **Table 2**), and List accounted for the known variation among lists of sentences within each test[1,2]. In cases where random effects had a small number of levels (e.g., Test, which had two levels), we follow recommendations by Oberpiller and colleagues[3] to treat them as random effects unless a singular fit occurs. These initial models did not include random slopes and are specified as “Design-constrained” in the **Supplementary Tables** below. Additional models were then built by iteratively adding or removing intercepts and slopes to the random effects structure following rules described in prior recommendations[4]. All candidate models, including final models, are reported in the **Supplementary Tables** below following best practice guidelines[5].

**Assessing order effects.** We additionally explored whether condition order (i.e., receiving off-target vs tcVNS first) impacted tcVNS efficacy. To this end, a two-level categorical variable (placebo-first, tcVNS-first) was included as a predictor to assess its interaction with Site (see **Supplementary Analyses**).

### Supplementary Tables

The tables below list the candidate models and estimates from the final model.

**Candidate models**. All tables with candidate models are ordered from most-to-least complex based on their degrees of freedom. Initial models that were constructed based on the experimental and stimulus design are highlighted in gray and specified as “Design-constrained”. AICc denotes the corrected Akaike Information Criterion, BIC the Bayesian Information Criterion, and LL the log-likelihood. Final models were selected on the basis of BIC and are highlighted in green. In cases where the final model is the design-constrained model, only green shading is shown.

**Final model estimates**. Tables containing the final model estimates also contain the model formula in its header.

**Table S1**. **Candidate models evaluating the impact of stimulation site on SNR-50**.

| **QuickSIN** | | | | | |
|---|---|---|---|---|---|
| Sampling units | Total obs = 172<br>Sites = 2; Subjects = 15; List = 12 | | | | |
| **Model specification** | **Model Formula** | **AICc** | **BIC** | **LL** | **df** |
| Random slopes & 2 intercepts | SNR50 ~ Site + (1+Site\|Subj) + (1+Site\|List) | Singular fit | | | 9 |
| Random slopes & 1 intercept | SNR50 ~ Site + (1+Site\|Subj) | Singular fit | | | 6 |
| Design-constrained | SNR50 ~ Site + (1\|Subj) + (1\|List) | 782.19 | 797.56 | -385.91 | 5 |
| Random intercept | SNR50 ~ Site + (1\|Subj) | 781.55 | 793.90 | -386.66 | 4 |
| No random effects | SNR50 ~ Site | 968.01 | 977.31 | -480.94 | 3 |
| **AzBio** | | | | | |
| Sampling units | Total obs = 28<br>Groups: Subjects = 14; List = 1 | | | | |
| **Model specification** | **Model Formula** | **AICc** | **BIC** | **LL** | **df** |
| Random slope & intercept | SNR50 ~ Site + (1+Site\|Subj) | Convergence failure | | | 6 |
| Design-constrained | SNR50 ~ Site + (1\|Subj) | 105.65 | 109.24 | -47.95 | 4 |
| No random effects | SNR50 ~ Site | 146.70 | 149.69 | -69.85 | 3 |

**Table S2. Final model estimates for tcVNS effects on SNR-50.**

| **QuickSIN:** SNR50 ~ Site + (1\|Subj) | | | | | |
|---|---|---|---|---|---|
| **Fixed Effects** | | | | | |
| | **Estimate** | **SE** | **95% CI** | **t** | **p** |
| Intercept | 6.26 | 0.91 | 4.43, 8.09 | 6.90 | 0.000015 |
| Site | -0.76 | 0.30 | -1.34, -0.17 | -2.53 | 0.016 |
| **Random Effects** | | | | | |
| | **Variance** | **SD** | | | |
| Subj (Intercept) | 12.46 | 3.53 | | | |
| Residual | 3.83 | 1.96 | | | |
| **Model Fit** | | | | | |
| $R^2$ | **Marginal** | **Conditional** | | | |
| | 0.0087 | 0.77 | | | |
| **AzBio:** SNR50 ~ Site + (1\|Subj) | | | | | |
| **Fixed Effects** | | | | | |
| | **Estimate** | **SE** | **95% CI** | **t** | **p** |
| Intercept | 2.83 | 0.81 | 1.18, 4.47 | 3.48 | 0.0079 |
| Site | -0.38 | 0.17 | -0.72, -0.033 | -2.22 | 0.045 |
| **Random Effects** | | | | | |
| | **Variance** | **SD** | | | |
| Subj (Intercept) | 9.06 | 3.01 | | | |
| Residual | 0.20 | 0.45 | | | |
| **Model Fit** | | | | | |
| $R^2$ | **Marginal** | **Conditional** | | | |
| | 0.0039 | 0.98 | | | |

**Table S3. Candidate models evaluating sex-related differences in tcVNS efficacy on SNR-50.**

| Sampling units | Total obs = 200<br>Sites = 2; Subjects = 22; List = 13; Test = 2; Sex = 2 | | | | |
|---|---|---|---|---|---|
| **Model specification** | **Model Formula** | **AICc** | **BIC** | **LL** | **df** |
| Random slopes & 2 intercepts, with nesting | SNR50 ~ Site*Sex + (1+Site\|Subj) + (1+Site\|Test/List) | Singular fit | | | 14 |
| Random slopes & 2 intercepts, without nesting | SNR50 ~ Site*Sex + (1+Site\|Subj) + (1+Site\|Test) | Singular fit | | | 11 |
| Random slopes & 2 intercepts | SNR50 ~ Site*Sex + (1+Site\|Subj) + (1+Site\|List) | Singular fit | | | 11 |
| Random slopes & nested intercept | SNR50 ~ Site*Sex + (1+Site\|Test/List) | Singular fit | | | 11 |
| Random slopes & intercept (for Subj) | SNR50 ~ Site*Sex + (1+Site\|Subj) | Singular fit | | | 8 |
| Random slopes & intercept (for Test) | SNR50 ~ Site*Sex + (1+Site\|Test) | Singular fit | | | 8 |
| Random slopes & intercept (for List) | SNR50 ~ Site*Sex + (1+Site\|List) | Singular fit | | | 8 |
| Design-constrained | SNR50 ~ Site*Sex + (1\|Subj) + (1\|Test/List) | 903.01 | 928.64 | -443.13 | 8 |
| 2 random intercepts | SNR50 ~ Site*Sex + (1\|Subj) + (1\|Test) | 903.04 | 925.55 | -444.23 | 7 |
| 2 random intercepts | SNR50 ~ Site*Sex + (1\|Subj) + (1\|List) | 915.09 | 937.59 | -450.25 | 7 |
| 1 nested random intercept | SNR50 ~ Site*Sex + (1\|Test/List) | Singular fit | | | 7 |
| 1 random intercept (for Test) | SNR50 ~ Site*Sex + (1\|Test) | 1095.39 | 1114.75 | -541.48 | 6 |
| 1 random intercept (for List) | SNR50 ~ Site*Sex + (1\|List) | 1104.57 | 1123.93 | -546.07 | 6 |
| 1 random intercept (for Subj) | SNR50 ~ Site*Sex + (1\|Subj) | 966.09 | 985.45 | -476.83 | 6 |
| No random effects | SNR50 ~ Site*Sex | 1112.46 | 1128.64 | -551.07 | 5 |

**Table S4. Final model estimates for sex-related differences in tcVNS efficacy on SNR-50.**

| **Model:** SNR50 ~ Site*Sex + (1 \| Subj) + (1 \| Test) | | | | | |
|---|---|---|---|---|---|
| **Fixed Effects** | | | | | |
| | **Estimate** | **SE** | **95% CI** | **t** | **p** |
| Intercept | 3.69 | 2.45 | -2.16, 9.52 | 1.51 | 0.33 |
| Site | -0.72 | 0.33 | -1.36, -0.078 | -2.20 | 0.029 |
| Sex | 1.67 | 1.56 | -1.45, 4.76 | 1.07 | 0.30 |
| Site:Sex | 0.054 | 0.57 | -1.06, 1.17 | 0.094 | 0.93 |
| **Random Effects** | | | | | |
| | **Variance** | **SD** | | | |
| Subj (Intercept) | 10.57 | 3.25 | | | |
| Test (Intercept) | 10.39 | 3.22 | | | |
| Residual | 3.592 | 1.90 | | | |
| **Model Fit** | | | | | |
| $R^2$ | **Marginal** | **Conditional** | | | |
| | 0.030 | 0.86 | | | |

**Table S5. Candidate models evaluating the impact of stimulation site on psychometric function parameters (SNRα and λ).**

| **QuickSIN—SNRα** | | | | | |
|---|---|---|---|---|---|
| Sampling units | Total obs = 32<br>Sites = 2; Subjects = 16 | | | | |
| **Model specification** | **Model Formula** | **AICc** | **BIC** | **LL** | **df** |
| Random slopes & intercept (for Subj) | SNRα ~ Site + (1+Site\|Subj) | Convergence failure | | | 6 |
| Design-constrained | SNRα ~ Site + (1\|Subj) | 126.86 | 131.24 | -58.69 | 4 |
| No random effects | SNR50 ~ Site | 137.46 | 141.00 | -65.30 | 3 |
| **QuickSIN—λ** | | | | | |
| Sampling units | Total obs = 32<br>Sites = 2; Subjects = 16 | | | | |
| **Model specification** | **Model Formula** | **AICc** | **BIC** | **LL** | **df** |
| Random slopes & intercept (for Subj) | λ ~ Site + (1+Site\|Subj) | Convergence failure | | | 6 |
| Design-constrained | λ ~ Site + (1\|Subj) | 220.97 | 225.35 | -105.74 | 4 |
| No random effects | λ ~ Site | 244.28 | 247.82 | -118.71 | 3 |
| **AzBio—SNRα** | | | | | |
| Sampling units | Total obs = 28<br>Sites = 2; Subjects = 14 | | | | |
| **Model specification** | **Model Formula** | **AICc** | **BIC** | **LL** | **df** |
| Random slopes & intercept (for Subj) | SNRα ~ Site + (1+Site\|Subj) | Convergence failure | | | 6 |
| Design-constrained | SNRα ~ Site + (1\|Subj) | 93.78 | 97.37 | -42.02 | 4 |
| No random effects | SNR50 ~ Site | 103.48 | 106.47 | -48.24 | 3 |
| **AzBio—λ** | | | | | |
| Sampling units | Total obs = 28<br>Sites = 2; Subjects = 14 | | | | |
| **Model specification** | **Model Formula** | **AICc** | **BIC** | **LL** | **df** |
| Random slopes & intercept (for Subj) | λ ~ Site + (1+Site\|Subj) | Convergence failure | | | 6 |
| Design-constrained | λ ~ Site + (1\|Subj) | 190.46 | 194.05 | -90.36 | 4 |
| No random effects | λ ~ Site | 230.36 | 233.35 | -111.68 | 3 |

**Table S6. Final model estimates for tcVNS effects on psychometric function parameters.**

| **QuickSIN:** SNRα ~ Site + (1 \| Subj) | | | | | |
|---|---|---|---|---|---|
| **Fixed Effects** | | | | | |
| | **Estimate** | **SE** | **95% CI** | **t** | **p** |
| Intercept | 3.71 | 0.48 | 2.75, 4.67 | 7.72 | $7.15 \times 10^{-7}$ |
| Site | -0.022 | 0.34 | -0.70, 0.66 | -0.066 | 0.95 |
| **Random Effects** | | | | | |
| | **Variance** | **SD** | | | |
| Subj (Intercept) | 2.78 | 1.67 | | | |
| Residual | 0.91 | 0.96 | | | |
| **Model Fit** | | | | | |
| **$R^2$** | **Marginal** | **Conditional** | | | |
| | 0.000035 | 0.75 | | | |

| **QuickSIN:** λ ~ Site + (1 \| Subj) | | | | | |
|---|---|---|---|---|---|
| **Fixed Effects** | | | | | |
| | **Estimate** | **SE** | **95% CI** | **t** | **p** |
| Intercept | 88.65 | 2.55 | 83.54, 93.76 | 34.74 | $1.02 \times 10^{-16}$ |
| Site | 3.74 | 1.43 | 0.86, 6.63 | 2.62 | 0.031 |
| **Random Effects** | | | | | |
| | **Variance** | **SD** | | | |
| Subj (Intercept) | 87.78 | 9.37 | | | |
| Residual | 16.39 | 4.05 | | | |
| **Model Fit** | | | | | |
| $R^2$ | **Marginal** | **Conditional** | | | |
| | 0.034 | 0.85 | | | |

| **AzBio:** SNRα ~ Site + (1 \| Subj) | | | | | |
|---|---|---|---|---|---|
| **Fixed Effects** | | | | | |
| | **Estimate** | **SE** | **95% CI** | **t** | **p** |
| Intercept | 1.10 | 0.38 | 0.35, 1.86 | 2.93 | 0.019 |
| Site | 0.037 | 0.24 | -0.46, 0.53 | 0.15 | 0.95 |
| **Random Effects** | | | | | |
| | **Variance** | **SD** | | | |
| Subj (Intercept) | 1.56 | 1.25 | | | |
| Residual | 0.41 | 0.64 | | | |
| **Model Fit** | | | | | |
| $R^2$ | **Marginal** | **Conditional** | | | |
| | 0.00018 | 0.79 | | | |

| **AzBio:** λ ~ Site + (1 \| Subj) | | | | | |
|---|---|---|---|---|---|
| **Fixed Effects** | | | | | |
| | **Estimate** | **SE** | **95% CI** | **t** | **p** |
| Intercept | 88.64 | 3.62 | 81.31, 95.96 | 24.47 | $5.68 \times 10^{-12}$ |
| Site | 2.43 | 0.99 | 0.42, 4.44 | 2.45 | 0.039 |
| **Random Effects** | | | | | |
| | **Variance** | **SD** | | | |
| Subj (Intercept) | 176.79 | 13.30 | | | |
| Residual | 6.91 | 2.63 | | | |
| **Model Fit** | | | | | |
| $R^2$ | **Marginal** | **Conditional** | | | |
| | 0.0083 | 0.96 | | | |

**Table S7. Candidate models evaluating the interaction of Site and SNR word recognition accuracy.**

| **QuickSIN** | | | | | |
|---|---|---|---|---|---|
| Sampling units | Total obs = 1032<br>Sites = 2; Subjects = 15; SNR = 6 | | | | |
| **Model specification** | **Model Formula** | **AICc** | **BIC** | **LL** | **df** |
| 3 random slopes & 1 random intercept | Acc ~ Site*SNR + (1+Site+SNR+Site:SNR\|Subj) | Singular fit | | | 14 |
| 2 random slopes & 1 random intercept | Acc ~ Site*SNR + (1+Site+Site:SNR\|Subj) | Convergence failure | | | 14 |
| 1 random slope (interaction) & 1 random intercept | Acc ~ Site*SNR + (1+Site:SNR\|Subj) | Singular fit | | | 10 |
| 2 random slopes & 1 random intercept | Acc ~ Site*SNR + (1+Site+SNR\|Subj) | Convergence failure | | | 10 |
| 2 random slopes & 1 random intercept | Acc ~ Site*SNR + (1+SNR+Site:SNR\|Subj) | Singular fit | | | 10 |
| Random slope & intercept | Acc ~ Site*SNR + (1+Site\|Subj) | Singular fit | | | 7 |
| Random slope & intercept | Acc ~ Site*SNR + (1+SNR\|Subj) | 2517.64 | 2552.11 | -1251.77 | 7 |
| Design-constrained | Acc ~ Site*SNR + (1\|Subj) | 2615.30 | 2639.94 | -1302.62 | 5 |
| No random effects | Acc ~ Site*SNR | 3036.03 | 3055.75 | -1514.00 | 4 |
| **AzBio** | | | | | |
| Sampling units | Total obs = 196<br>Groups: Subjects = 14; List = 1 | | | | |
| **Model specification** | **Model Formula** | **AICc** | **BIC** | **LL** | **df** |
| 3 random slopes & 1 random intercept | Acc ~ Site*SNR + (1+Site+SNR+Site:SNR\|Subj) | Singular fit | | | 14 |
| 2 random slopes & 1 random intercept | Acc ~ Site*SNR + (1+Site+Site:SNR\|Subj) | Singular fit | | | 14 |
| 1 random slope (interaction) & 1 random intercept | Acc ~ Site*SNR + (1+Site:SNR\|Subj) | Singular fit | | | 10 |
| 2 random slopes & 1 random intercept | Acc ~ Site*SNR + (1+Site+SNR\|Subj) | 1810.03 | 1841.62 | -894.42 | 10 |
| 2 random slopes & 1 random intercept | Acc ~ Site*SNR + (1+SNR+Site:SNR\|Subj) | Singular fit | | | 10 |
| Random slope & intercept | Acc ~ Site*SNR + (1+Site\|Subj) | 2610.37 | 2632.72 | -1297.89 | 7 |
| Random slope & intercept | Acc ~ Site*SNR + (1+SNR\|Subj) | 1818.24 | 1840.59 | -901.82 | 7 |
| Design-constrained | Acc ~ Site*SNR + (1\|Subj) | 2614.45 | 2630.52 | -1302.07 | 5 |
| No random effects | Acc ~ Site*SNR | 5651.08 | 5663.97 | -2821.44 | 4 |

**Table S8. Final model estimates for the interaction between tcVNS effects and SNR on SIN intelligibility**

| **QuickSIN:** Acc ~ Site*SNR + (1 + SNR \| Subj) | | | | | |
|---|---|---|---|---|---|
| **Fixed Effects** | | | | | |
| | **Estimate** | **SE** | **95% CI** | **z** | **p** |
| Intercept | -1.07 | 0.133 | -1.34, -0.80 | -8.02 | $2.78 \times 10^{-15}$ |
| Site | -0.016 | 0.12 | -0.25, 0.22 | -0.13 | 0.90 |
| SNR | 0.23 | 0.020 | 0.19, 0.27 | 11.49 | $6.11 \times 10^{-30}$ |
| Site:SNR | 0.028 | 0.011 | 0.0052, 0.050 | 2.42 | 0.021 |
| **Random Effects** | | | | | |
| | **Variance** | **SD** | **Corr** | | |
| Subj (Intercept) | 0.16 | 0.41 | | | |
| SNR | 0.0048 | 0.069 | 0.33 | | |
| **Model Fit** | | | | | |
| **$R^2$** | **Marginal** | **Conditional** | | | |
| | 0.67 | 0.89 | | | |
| **AzBio:** Acc ~ Site*SNR + (1 + SNR \| Subj) | | | | | |
| **Fixed Effects** | | | | | |
| | **Estimate** | **SE** | **95% CI** | **z** | **p** |
| Intercept | -0.58 | 0.15 | -0.90, -0.27 | -3.87 | 0.00022 |
| Site | 0.039 | 0.044 | -0.048, 0.13 | 0.88 | 0.43 |
| SNR | 0.32 | 0.28 | 0.26, 0.38 | 11.49 | $6.11 \times 10^{-30}$ |
| Site:SNR | 0.025 | 0.0067 | 0.012, 0.039 | 3.81 | 0.00023 |
| **Random Effects** | | | | | |
| | **Variance** | **SD** | **Corr** | | |
| Subj (Intercept) | 0.30 | 0.55 | | | |
| SNR | 0.010 | 0.10 | 0.85 | | |
| **Model Fit** | | | | | |
| **$R^2$** | **Marginal** | **Conditional** | | | |
| | 0.73 | 0.99 | | | |

**Table S9. Candidate models evaluating established relationships between hearing loss (PTA4), baseline speech-in-noise ability (SNR-50), word recognition accuracy (Acc), psychometric function parameters (SNRα and λ), and Age.**

| **PTA4 vs SNR50** | | | | | |
|---|---|---|---|---|---|
| Sampling units | Total obs = 30<br>Test = 2 | | | | |
| **Model specification** | **Model Formula** | **AICc** | **BIC** | **LL** | **df** |
| Design-constrained | SNR50 ~ PTA4 + (1\|Test) | 166.44 | 170.45 | -78.42 | 4 |
| No random effects | SNR50 ~ PTA4 | 164.08 | 167.36 | -78.58 | 3 |
| **PTA4 vs Proportion Correct** | | | | | |
| Sampling units | Total obs = 30<br>Test = 2 | | | | |
| **Model specification** | **Model Formula** | **AICc** | **BIC** | **LL** | **df** |
| Design-constrained | PC ~ PTA4 + (1\|Test) | 240.72 | 244.73 | -115.56 | 4 |
| No random effects | PC ~ PTA4 | 241.31 | 244.59 | -117.19 | 3 |
| **PTA4 vs SNRα** | | | | | |
| Sampling units | Total obs = 30<br>Test = 2 | | | | |
| **Model specification** | **Model Formula** | **AICc** | **BIC** | **LL** | **df** |
| Design-constrained | SNRα ~ PTA4 + (1\|Test) | 114.11 | 118.12 | -52.26 | 4 |
| No random effects | SNRα ~ PTA4 | 121.17 | 124.45 | -57.12 | 3 |
| **PTA4 vs λ** | | | | | |
| Sampling units | Total obs = 30<br>Test = 2 | | | | |
| **Model specification** | **Model Formula** | **AICc** | **BIC** | **LL** | **df** |
| Design-constrained | λ ~ PTA4 + (1\|Test) | 242.34 | 246.35 | -116.37 | 4 |
| No random effects | λ ~ PTA4 | 242.06 | 245.34 | -117.57 | 3 |
| **Age vs SNR50** | | | | | |
| Sampling units | Total obs = 30<br>Test = 2 | | | | |
| **Model specification** | **Model Formula** | **AICc** | **BIC** | **LL** | **df** |
| Design-constrained | SNR50 ~ Age + (1\|Test/List) | 579.78 | 592.17 | -284.57 | 5 |
| Remove nesting, List | SNR50 ~ Age + (1\|Test) | 577.56 | 587.56 | -284.57 | 4 |
| No random effects | SNR50 ~ Age | 576.41 | 583.98 | -285.08 | 3 |

**Table S10. Final model estimates for each established relationship.**

| **PTA4 vs SNR50:** SNR50 ~ PTA4 | | | | | |
|---|---|---|---|---|---|
| **Fixed Effects** | | | | | |
| | **Estimate** | **SE** | **95% CI** | **t** | **p** |
| Intercept | -0.23 | 1.73 | -3.77, 3.31 | -0.13 | 0.98 |
| PTA4 | 0.20 | 0.064 | 0.064, 0.33 | 3.04 | 0.012 |
| **Model Fit** | | | | | |
| **$R^2$** | **Multiple** | **Adjusted** | | | |
| | 0.25 | 0.22 | | | |
| **PTA4 vs Proportion Correct:** PC ~ PTA4 | | | | | |
| **Fixed Effects** | | | | | |
| | **Estimate** | **SE** | **95% CI** | **t** | **p** |
| Intercept | 83.12 | 6.87 | 70.27, 95.98 | 12.10 | $1.36 \times 10^{-5}$ |

| | | | | | |
|---|---|---|---|---|---|
| PTA4 | -0.62 | 0.23 | -1.05, -0.14 | -2.72 | 0.022 |
| **Random Effects** | | | | | |
| | **Variance** | **SD** | | | |
| Test (Intercept) | 21.23 | 4.61 | | | |
| Residual | 144.01 | 12.00 | | | |
| **Model Fit** | | | | | |
| **$R^2$** | **Marginal** | **Conditional** | | | |
| | 0.18 | 0.29 | | | |
| **PTA4 vs SNRα:** SNRα ~ PTA4 + (1 \| Test) | | | | | |
| **Fixed Effects** | | | | | |
| | **Estimate** | **SE** | **95% CI** | **t** | **p** |
| Intercept | -0.032 | 1.30 | -2.94, 2.89 | -0.024 | 0.98 |
| PTA4 | 0.098 | 0.023 | 0.053, 0.14 | 4.31 | 0.00077 |
| **Random Effects** | | | | | |
| | **Variance** | **SD** | | | |
| Test (Intercept) | 2.64 | 1.62 | | | |
| Residual | 1.45 | 1.20 | | | |
| **Model Fit** | | | | | |
| **$R^2$** | **Marginal** | **Conditional** | | | |
| | 0.19 | 0.71 | | | |
| **PTA4 vs λ:** λ ~ PTA4 | | | | | |
| **Fixed Effects** | | | | | |
| | **Estimate** | **SE** | **95% CI** | **t** | **p** |
| Intercept | 99.35 | 6.33 | 86.38, 112.32 | 15.69 | $2.53 \times 10^{-14}$ |
| PTA4 | -0.43 | 0.24 | -0.91, 0.055 | -1.82 | 0.11 |
| **Model Fit** | | | | | |
| **$R^2$** | **Multiple** | **Adjusted** | | | |
| | 0.11 | 0.073 | | | |
| **Age vs SNR50:** SNR50 ~ Age | | | | | |
| **Fixed Effects** | | | | | |
| | **Estimate** | **SE** | **95% CI** | **t** | **p** |
| Intercept | -4.46 | 4.40 | -13.19, 4.27 | -1.01 | 0.38 |
| Age | 0.15 | 0.062 | 0.023, 0.27 | 2.37 | 0.034 |
| **Model Fit** | | | | | |
| **$R^2$** | **Multiple** | **Adjusted** | | | |
| | 0.054 | 0.044 | | | |
| **Age vs PTA4:** PTA4 ~ Age | | | | | |
| **Fixed Effects** | | | | | |
| | **Estimate** | **SE** | **95% CI** | **t** | **p** |
| Intercept | -29.9 | 15.2 | -61.02, 1.25 | -1.97 | 0.0593 |
| Age | 0.78 | 0.22 | 0.34, 1.22 | 3.63 | 0.0011 |
| **Model Fit** | | | | | |
| **$R^2$** | **Multiple** | **Adjusted** | | | |
| | 0.32 | 0.29 | | | |

**Table S11. Variance Inflation Factors for each candidate multivariable mixed-effects model.**

| **Model:** Effect ~ SNR50 * PTA4 * Age + (1\|Test) | | |
|---|---|---|
| | **VIF** | **VIF 95% CI** |
| SNR50 | 2.06 | 1.53, 3.15 |
| PTA4 | 4.75 | 3.18, 7.45 |
| Age | 2.15 | 1.58, 3.28 |
| SNR50:PTA4 | 1.99 | 1.48, 3.03 |
| SNR50:Age | 3.24 | 2.25, 5.01 |
| PTA4:Age | 2.09 | 1.54, 3.19 |
| SNR50:PTA4:Age | 4.44 | 2.99, 6.95 |
| **Model:** Effect ~ SNR50 * PTA4 * Age * PropCorrect + (1\|Test) | | |
| SNR50 | 1399.58 | 1041.28, 1881.28 |
| PTA4 | 15.99 | 12.04, 21.35 |
| Age | 6.71 | 5.14, 8.88 |
| PC | 1370.85 | 1019.91, 1842.67 |
| SNR50:PTA4 | 7514.15 | 5589.86, 10100.97 |
| SNR50:Age | 259.54 | 193.22, 348.75 |
| PTA4:Age | 31.61 | 23.66, 42.35 |
| SNR50:PC | 13.13 | 9.91, 17.50 |
| PTA4:PC | 7162.58 | 5328.34, 9628.37 |
| Age:PC | 261.92 | 194.99, 351.95 |
| SNR50:PTA4:Age | 3220.25 | 2395.66, 4328.77 |
| SNR50:PTA4:PC | 46.90 | 35.03, 62.90 |
| SNR50:Age:PC | 16.59 | 12.49, 22.15 |
| PTA4:Age:PC | 3289.39 | 2447.10, 4421.72 |
| SNR50:PTA4:Age:PC | 42.04 | 31.42, 56.36 |
| **Model:** Effect ~ SNR50 * PTA4 * Age * MoCA + (1\|Test) | | |
| SNR50 | 24.51 | 18.38, 32.81 |
| PTA4 | 15.91 | 11.98, 21.25 |
| Age | 11.03 | 8.35, 14.68 |
| MoCA | 14.33 | 10.81, 19.12 |
| SNR50:PTA4 | 29.73 | 22.26, 39.82 |
| SNR50:Age | 12.65 | 9.56, 16.87 |
| PTA4:Age | 77.51 | 57.81, 104.06 |
| SNR50:MoCA | 19.36 | 14.55, 25.88 |
| PTA4:MoCA | 15.50 | 11.68, 20.69 |
| Age:MoCA | 12.34 | 9.33, 16.44 |
| SNR50:PTA4:Age | 191.39 | 142.52, 257.13 |
| SNR50:PTA4:MoCA | 30.72 | 23.00, 41.15 |
| SNR50:Age:MoCA | 17.89 | 13.45, 23.90 |
| PTA4:Age:MoCA | 103.61 | 77.22, 139.14 |
| SNR50:PTA4:Age:MoCA | 174.21 | 129.74, 234.04 |

**Table S12. Candidate models evaluating the relationship between baseline measures (SNR-50, Age, and PTA4) and the magnitude of tcVNS-evoked effects.**

| Sampling units | Total obs = 30<br>Sites = 2; Test = 2 | | | | |
|---|---|---|---|---|---|
| **Model specification** | **Model Formula** | **AICc** | **BIC** | **LL** | **df** |
| Design-constrained | Effect ~ SNR50*PTA4*Age + (1\|Test) | 173.98 | 176.41 | -71.20 | 10 |
| No random effects | Effect ~ SNR50*PTA4*Age | 182.07 | 185.68 | -77.54 | 9 |

**Table S13. Final model estimates for the relationship between baseline measures and the magnitude of tcVNS-evoked effects.**

| **Model:** Effect ~ SNR50*PTA4*Age + (1 \| Test) | | | | | |
|---|---|---|---|---|---|
| **Fixed Effects** | | | | | |
| | **Estimate** | **SE** | **95% CI** | **t** | **p** |
| Intercept | 3.34 | 0.95 | 1.28, 5.24 | 3.53 | 0.10 |
| SNR50 | 3.59 | 1.01 | 1.78, 5.30 | 3.55 | 0.028 |
| PTA4 | -0.96 | 1.51 | -3.58, 1.70 | -0.63 | 0.97 |
| Age | 0.91 | 1.01 | -0.84, 2.70 | 0.89 | 0.88 |
| SNR50:PTA4 | -0.43 | 0.82 | -1.83, 1.04 | -0.52 | 0.99 |
| SNR50:Age | 1.58 | 2,03 | -1.94, 5.15 | 0.78 | 0.93 |
| PTA4:Age | -0.73 | 1.13 | -2.77, 1.18 | -0.64 | 0.97 |
| SNR50:PTA4:Age | -0.33 | 1.63 | -3.14, 2.56 | -0.21 | 1.00 |
| **Random Effects** | | | | | |
| | **Variance** | **SD** | | | |
| Test (Intercept) | 0.37 | 0.61 | | | |
| Residual | 13.81 | 3.72 | | | |
| **Model Fit** | | | | | |
| $R^2$ | **Marginal** | **Conditional** | | | |
| | 0.48 | 0.49 | | | |

**Table S14. Final model estimates for the interaction of tcVNS and the order of stimulation.**

| **QuickSIN:** SNR50 ~ Site*Order + (1 \| Subj) | | | | | |
|---|---|---|---|---|---|
| **Fixed Effects** | | | | | |
| | **Estimate** | **SE** | **95% CI** | **t** | **p** |
| Intercept | 5.70 | 1.31 | 3.15, 8.25 | 4.35 | 0.00059 |
| Site | -0.74 | 0.42 | -1.57, 0.086 | -1.76 | 0.081 |
| Order | 1.12 | 1.85 | -2.48, 4.73 | 0.61 | 0.55 |
| Site:Order | -0.023 | 0.60 | -1.20, 1.15 | -0.039 | 0.97 |
| **Random Effects** | | | | | |
| | **Variance** | **SD** | | | |
| Subj (Intercept) | 13.02 | 3.61 | | | |
| Residual | 3.86 | 1.96 | | | |
| **Model Fit** | | | | | |
| $R^2$ | **Marginal** | **Conditional** | | | |
| | 0.026 | 0.78 | | | |
| **AzBio:** SNR50 ~ Site*Order + (1 \| Subj) | | | | | |
| **Fixed Effects** | | | | | |
| | **Estimate** | **SE** | **95% CI** | **t** | **p** |
| Intercept | 2.48 | 1.19 | 0.17, 4.80 | 2.09 | 0.059 |
| Site | -0.31 | 0.25 | -0.79, 0.17 | -1.24 | 0.24 |
| Order | 0.69 | 1.68 | -2.58, 3.96 | 0.41 | 0.69 |
| Site:Order | -0.14 | 0.35 | -0.82, 0.54 | -0.40 | 0.79 |
| **Random Effects** | | | | | |
| | **Variance** | **SD** | | | |
| Subj (Intercept) | 9.70 | 3.11 | | | |
| Residual | 0.21 | 0.46 | | | |
| **Model Fit** | | | | | |
| $R^2$ | **Marginal** | **Conditional** | | | |
| | 0.014 | 0.98 | | | |

### Supplementary Analyses

#### SIN intelligibility and hearing loss followed established patterns

To ensure consistency with prior research, we assessed several relationships that are expected to exist between baseline SIN performance (i.e., during off-target/placebo stimulation), hearing loss (PTA4), and age. Model candidates are listed in **Table S9** and final LME model estimates are shown in **Table S10**. First, we replicated the link between hearing loss and SIN intelligibility[6-13]. Both SNR-50 (**Figure S1A**; p=0.012) and accuracy (**Figure S1B**; p=0.022) diminished with hearing loss. Second, we demonstrated the hypothesized relationship between the psychometric function's midpoint (SNRα) and hearing losss[11]: SNRα increased with PTA4 (**Figure S1C**; p=0.00077). In contrast, the upper asymptote (λ) did not vary with PTA4 (**Figure S1D**; p=0.11), supporting the hypothesis that λ reflects independent contributions by central auditory processing and cognitive function[11]. Third, baseline SNR-50 increased with age (**Figure S1E**; p=0.034), corroborating prior research[8,10,11,14]. Similarly, hearing loss increased with age[15] (**Figure S1F**; p=0.0011).

#### Non-significant effect of stimulation order on tcVNS efficacy

To evaluate whether the order of conditions (tcVNS and off-target stimulation) influenced the results, we extended the final model formula that assessed effects of tcVNS on SNR-50 (**Table S1**) to include a categorical variable for Order (2 levels: placebo-first, tcVNS-first; **Table S14**). There was an absence of significant interaction effects between Site and Order, indicating that the order in which conditions were presented did not systematically influence tcVNS-evoked changes. However, we note that adding an additional predictor to the model can lead to a loss of statistical power, which could obscure a small, but present effect of Order.

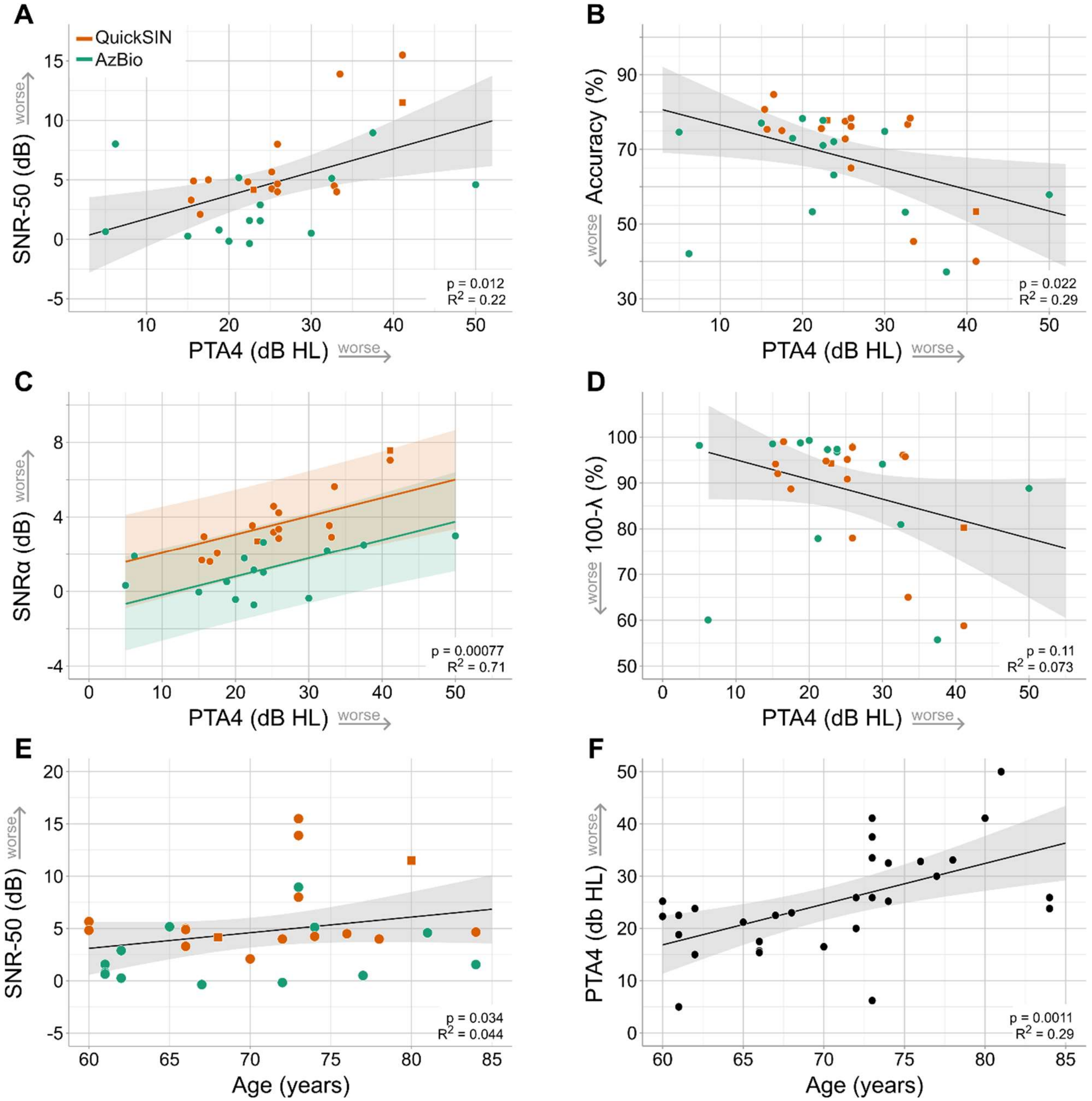

**Figure S1. Speech-in-noise intelligibility and hearing loss followed established patterns.** Baseline speech-in noise metrics, measured during off-target stimulation, are displayed as a function of hearing loss (PTA4): (**A**) SNR-50, (**B**) word recognition accuracy, (**C**) SNRα, and (**D**) λ. (**E**) Baseline SNR-50 and (**F**) hearing loss is displayed as a function of participant age. In all plots, each dot depicts individual participants in a test cohort, lines show the best-fitting regression determined using models in Table S10, with shaded areas indicating 95% confidence intervals. All p-values are corrected for multiple comparisons using the Benjamini-Hochberg procedure[16].
